# Visualization of syntrophic benzene-fermenting *Desulfobacterota* ORM2 in a methanogenic enrichment culture using fluorescence *in situ* hybridization

**DOI:** 10.1101/2023.07.16.549207

**Authors:** Xu Chen, Courtney R.A. Toth, Shen Guo, Fei Luo, Jane Howe, Camilla L. Nesbø, Elizabeth A. Edwards

## Abstract

Although benzene degradation under strictly anoxic conditions was first reported over 25 years ago, the mechanism for benzene activation in the absence of oxygen is still elusive. A major limitation has been the difficulty to grow anaerobic benzene-degrading enrichment cultures. Our laboratory has maintained a methanogenic enrichment culture for decades, harboring a benzene fermenter referred to as *Desulfobacterota* ORM2. Recent genomic analyses indicate that ORM2 is not affiliated with any characterized genus, but it is phylogenetically similar to several other known and predicted benzene degraders. *Desulfobacterota* ORM2 has a doubling time of approximately 30 days and often enters a long lag or decay phase after inoculation into sterile pre-reduced anaerobic medium. A specific fluorescent *in situ* hybridization (FISH) probe was used to observe *Desulfobacterota* ORM2 cells during this decay phase, revealing a rod-shaped cell of variable length with a tendency to associate with other cells, particularly methanogens. Microscopic and genomic analyses indicate that *Desulfobacterota* ORM2 may produce extracellular polymeric substances (EPS) that likely contribute to cell aggregation. The production of EPS may consume a significant amount of energy, perhaps contributing to the lag time before onset of growth of *Desulfobacterota* ORM2 post-inoculation. We observed little cell aggregation in a culture amended with very high concentrations of benzene (90-120 mg/L). This study visualized the cells of a novel clade within the *Desulfobacterota* for the first time, enabling monitoring of spatial organization within a methanogenic consortium and provides hints to improve the growth rate of ORM2.

**Importance:** A specific FISH probe was designed for the poorly characterized benzene fermenter *Desulfobacterota* ORM2. This probe was used to monitor changes in spatial organization in a methanogenic benzene-degrading enrichment culture. ORM2 cells were often found in cell aggregates, revealing a possible reason for the long lag phases observed after inoculation.

## Introduction

Benzene is a widespread groundwater contaminant owing to the ubiquitous use of petroleum products and accidental releases from extraction, refining, and processing operations. Due to the carcinogenicity and low regulatory limits for benzene in groundwater, some form of remediation is normally required to accelerate nature attenuation at contaminated sites. Excavation and aeration are the most common methods for shallow spills due to rapid aerobic degradation.

However, for oxygen-limited contaminant plumes, such as deep aquifers and low permeability sites, or for inaccessible sites under buildings, anaerobic biodegradation processes contribute most to remediation. Anaerobic benzene biodegradation has been observed under iron-reducing, nitrate-reducing, sulfate-reducing, and methanogenic conditions (1–4). Methanogenic degradation is mostly found in regions depleted of all other electron acceptors, and often close to petroleum contaminant source zones (5). To promote remediation efforts, our laboratory is currently field testing bioaugmentation with a methanogenic benzene-degrading culture referred to as DGG-B^TM^. This culture is a large-scale (>100 L) sublineage of a longstanding (>20 years), laboratory-scale (<10 L) culture known as the “OR consortium” (1–3). The subculturing history of the “OR consortium” including the DGG-B^TM^ sublineage is shown in Figure S1. Growth studies and molecular analyses of OR and DGG-B^TM^ have identified a specialized benzene fermenter *Desulfobacterota* referred to as ORM2. Luo et al. (1) confirmed a direct correlation between anaerobic benzene degradation rates and 16S rRNA gene copy abundance of ORM2. Our laboratory has used enrichment culture experiments, 16S rRNA gene amplicon sequencing, metagenomic sequencing, and quantitative polymerase chain reactions (qPCR) to investigate methanogenic benzene degradation, as have a handful of other labs in with their own established methanogenic or sulfate-reducing benzene-degrading enrichment cultures (1, 2, 6-10). The microbial community composition of these cultures is remarkably similar, each harboring a key benzene degrading bacterium closely related to ORM2 (Figure 1). Microscopic investigations of the spatial distribution of cells and interspecies relationships, however, are rare due to the complexity of the microbial community. Fluorescence *in situ* hybridization (FISH) is a widely used technique to visualize target cells using rRNA-specific oligonucleotide probes. Developing such a tool for ORM2 is a logical next step for better characterization of the OR culture. Perhaps morphologies and spatial organization will provide clues to important interspecies interactions and community responses to changes in growth conditions (11).

**Figure 1.**
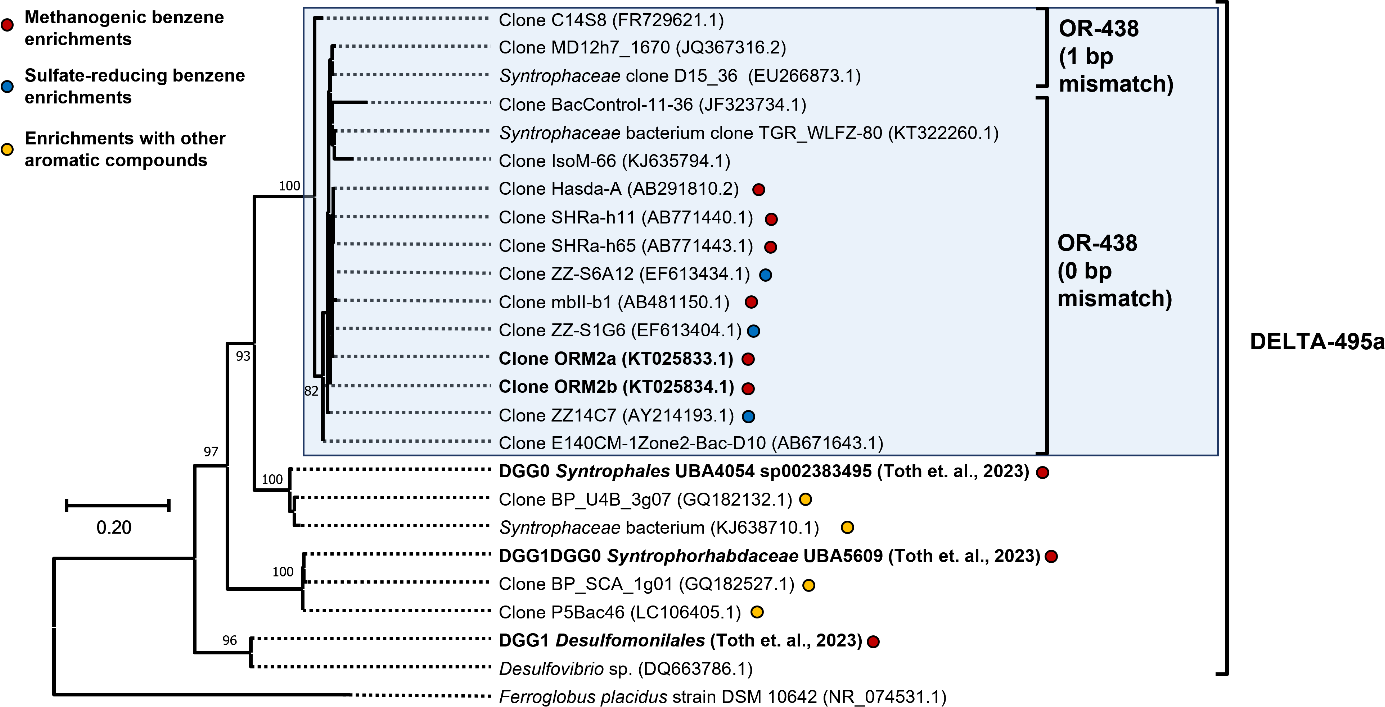
Maximum likelihood consensus tree showing affiliation of all the targets of probe OR-438 and selected strains targeted by probe DELTA-495a. Names in bold are microbes found in the OR/DGG consortium. All strains in the SILVA non-redundant 16S rRNA database, except for the strains highlighted in the boxed region of this figure, exhibit more than five mismatches. A description of how the final consensus tree was constructed is provided in SI (Text S1)

Maintaining methanogenic benzene degrading cultures is difficult because of persistent low cell densities, long doubling times (∼30 days), and long lag times after dilution of the culture into fresh medium. As a result, few laboratories maintain such cultures. A previous study from our lab by Luo et al. (1) monitored 1% and 0.1% transfers of an active benzene-degrading consortium over 415 days, where ORM2 numbers initially decreased during the first 50 days and long lag times (200 and 250 days, respectively) were observed before the onset of measurable benzene degradation. The abundance of benzene degraders (ORM2) was monitored by qPCR and benzene degradation was only detectable when ORM2 concentrations exceeded 1×10^5^ 16S rRNA copies/mL. A similar ORM2 concentration threshold for the onset of detectable benzene degradation was observed more recently by Toth et al. (2), but the reason for such long lag times was not determined. To better understand the community behavior and to investigate the reason for the long lag time, we designed and validated an ORM2-specific FISH probe. Next, we conducted a dilution and growth study where we monitored benzene degradation and microbial growth, collecting samples for FISH at multiple time points over ORM2’s growth cycle. This study revealed the morphology and spatial aggregation of these specialized benzene fermenters within a community of other microbes, particularly archaea. The ORM2 cell aggregates were observed to grow in both size and quantity over time while capturing other secondary fermenters and methanogenic archaea. As well, the aggregates showed a response to benzene concentration, with greater cell dispersion when benzene feeding concentrations were increased from 10 mg/L to 120 mg/L.

## Materials and Methods

### Chemicals and microbial cultures

All chemicals used in this study were obtained from Sigma-Aldrich (Oakville, ON, Canada) at the highest available purity, unless otherwise specified. The cultures were maintained in glass batch bottles ranging from 40 mL to 2 L at the University of Toronto and in large stainless steel vessels (20L-100L) at SiREM (Guelph, ON, Canada) following previously described protocols (3). Five methanogenic benzene-degrading culture bottles were examined: OR-b1A, DGG0, DGG4, DGG100 original and DGG100 pos B. The lineage relationship between these cultures is presented in Figure S1. At the time of sampling for microscopy and DNA extraction, OR-b1A, DGG4 and DGG100 original were fed with benzene to a target aqueous concentration of 20 mg/L, DGG0 to a target aqueous concentration of 40 mg/L, and DGG100 pos B to a target aqueous concentration of 120 mg/L.

### Culture maintenance and time-course growth experiment

This experiment was set up in 250 mL glass bottles sealed with screwcap Mininert valves. Bottles were set up in biological duplicates containing 40 mL of OR culture from parent culture bottle “DGG100 original” (see Figure S1) and 160 mL of iron sulfide (FeS)-reduced, bicarbonate-based mineral medium as previously described (3). Bottles were first purged for 20 mins to remove residual benzene and then fed with 2.5 µL of neat benzene (28 µmol/bottle), corresponding to an initial aqueous concentration of ∼10 mg/L. The duplicate bottles were fed regularly as shown in Table S1. Samples (1 mL for DNA extraction and 2 mL for FISH) were collected every week for the first 78 days. Afterwards, cultures were sampled on days 92, 111, 141, 188, 203, 245, and 300 for FISH and DNA (Table S1).

### Fluorescence *in situ* hybridization (FISH)

Probe Arch915 (12) was used to detect Archaea in this study. Probe DELTA-495a (21) was initially used to visualize most *Desulfobacterota* (previously *Deltaproteobacteria*). The design of specific FISH probes for ORM2 was conducted using the phylogeny software ARB (13). The 16S rRNA sequence of ORM2 was manually added into the SILVA non-redundant 16S rRNA database. All potential probes with GC content over 50% were selected and BLAST searched on NCBI to check for specificity. All probes which passed this initial screening test were analyzed using the oligonucleotide probe evaluation tool mathFISH (14–16) to find an optimal formamide concentration. All the FISH probes used in this study are summarized in Table S2a. Three ORM2-specific FISH probes were tested at respective optimal formamide concentrations and evaluated.

All samples for FISH were first harvested by centrifugation of a 2 mL culture sample at 13,000 x g for 20 mins. The supernatants were discarded, and the pellets were resuspended in 950 µL of 1x phosphate-buffered saline (PBS). Fifty microliters of a 20% paraformaldehyde (PFA) fixative solution were then added to achieve a 1% final concentration. The mixtures were then stored in a 4°C fridge for at least 24 hours to ensure complete fixation. After incubation, each mixture was centrifuged again (13,000 x g for 20 mins) and the resulting pellets were resuspended in 500 µL PBS twice to wash off the fixative. Aliquots (10 µL) of fixed cells were then loaded onto PTFE printed glass slides (cat#63429-04 from Electron Microscopy Sciences) with 4-mm diameter wells and allowed to air-dry. The air-dried glass slides were then dehydrated for 3 mins each in 50, 80, and 100% ethanol. Next, the slides were loaded with hybridization buffer (17) and incubated for two hours in a 46°C incubator. The hybridized samples were then incubated with washing buffer in a 48°C water bath as described previously (17). After washing, the slides were stained with 4’,6-diamidino-2-phenylindole (DAPI) and mounted in a 4:1 mixture of Citifluor (Citifluor Ltd, London, UK) and Vecta Shield (Vector Laboratories, Inc., Burlingame, CA), covered with glass coverslips, and sealed with nail polish.

### Epifluorescence Microscopy

A droplet of immersion oil was loaded onto the prepared FISH slides and the slides were observed using an epifluorescent microscope (BX 51, Olympus) with a 100x UPlan Apochromat objective, 150 W xenon lamp (Opti Quip), a set of excitation filter cubes (DAPI/Cy3/Cy5) and a 10x focusing eyepiece (1000-fold total magnification). The microscopic images were captured by a Hamamatsu ORCA-Flash 4.0 digital CMOS camera.

### DNA extraction, quantitative PCR, and Illumina amplicon sequencing

For every sample in this study, DNA was extracted from 1 mL of culture. Cells were harvested by centrifugation at 13,000 x g for 20 min at 4°C and pellets were resuspended in the remaining 0.05 mL supernatant. DNA was extracted using the MO BIO PowerSoil® DNA isolation kit following the manufacturer’s recommendations. Quantitative PCR assays were performed to track the 16S rRNA gene copy numbers of total Bacteria, total Archaea, and *Desulfobacterota* (ORM2), using primer pairs shown in Table S3. Amplicon sequencing of 16S rRNA genes was conducted using primers targeting the V6-V8 hypervariable region (Table S3). Amplification and sequencing were performed at the Centre d’expertise et de services Génome Québec (Montreal, Canada). Processing of 16S rRNA sequencing data into amplicon sequence variants (ASVs) was implemented using the DADA2 algorithm within version 2019.10 of QIIME2 (18), and classification of ASVs was performed using version 132 of the Silva SSU (small subunit rRNA, 16S/18S) database (19) (https://www.arb-silva.de/documentation/release-132/).

### Data availability

The closed genome sequence of *Desulfobacterota* ORM2 is available in the Joint Genome Institute (IMG) database under the GOLD ID Ga0311013 and in NCBI under the accession number CP113000.1 (20). Amplicon sequences (16S rRNA) generated in this study are available in the NCBI Short Read Archive (SRA) under the BioProject accession number PRJNA937673.

## Results and Discussion

### ORM2 cells are rods with variable lengths

The previously validated FISH probe DELTA-495a (21) was initially used as a positive control to visualize ORM2 in the methanogenic benzene-degrading culture. This broad specificity probe targets most *Desulfobacterota* including the main benzene fermenter ORM2. To verify the specificity of probe DELTA-495a in our culture, the probe’s nucleotide sequence was aligned against draft and complete (closed) metagenome-assembled genomes from two subcultures, DGG0 and DGG1A (20). In addition to ORM2, nucleotide BLAST results showed hits to three (draft) genomes from members of the *Desulfobacterota* (a *Syntrophales*, a *Syntrophorhabdaceae*, and a *Desulfomonilales*; Figure 1 and Table S4). However, the relative abundances of these three microbes are quite low (≤1.05%) based on metagenome coverage estimates. The DELTA-495a probe (Figure S2) was applied to the subculture OR-b1A, revealing rod-shaped microbes with variable cell lengths.

To eliminate possible interference from low abundance *Desulfobacterota* other than ORM2, three highly specific FISH probes for ORM2 were designed and tested (Table S2a). Only probes OR-436 and OR-438 resulted in a distinguishable fluorescence signal above the background noise. Probe OR-438 (18 base pairs, bp) showed higher fluorescence signal intensity than the longer probe OR-436 (20 bp), indicating a better hybridization efficiency. Therefore, OR-438 was selected for future experiments. Using the Probe Match tool with the Ribosomal Database Project (RDP), a specificity check of OR-438 returned only 16 hits with a maximum of 1 base pair mismatch (Table S2b). The 16 matches were all affiliated to a notable unclassified clade of *Desulfobacterota* that includes only strains from benzene-degrading enrichments (Figure 1). This clade includes ORM2 as well as the strains Hasda-A, SHRa-h11, SHRa-h65, and mbll-b1, which were recovered from methanogenic benzene-degrading enrichment cultures maintained by a Japanese research group (7). Further, the clade includes strains ZZ-14C7, ZZ-S1G6, and ZZ-S6A12 found in sulfate-reducing benzene enrichment cultures from a German research group (10), and strain IsoM-66 from a Canadian methanogenic oil sands tailing pond enrichment (22). Other matches are all from environmental samples that may contain benzene, including lake sediment, groundwater, or petroleum-contaminated plumes. A maximum likelihood consensus tree (Figure 1) shows all 16 hits clustering into a single clade. The taxonomic classification of this clade is currently under investigation; version 128 of the Silva database suggests that these organisms are affiliated with the candidate order Sva0485 (2), but more recent metagenomic insights indicate that this clade may instead represent a novel genus within the family *Syntrophaceae* (23). The specificity of probe OR-438 appears to make it a good biomarker for strictly anaerobic benzene-degrading microbes in environmental samples.

Example microscopic images of the OR-b1A subculture labelled with the OR-438 probe are shown in Figure 2. Although *Desulfobacterota* other than ORM2 were excluded with this more specific probe, the FISH images still revealed two different morphologies as shown in Figure 2, panels (d) and (e). One morphology is a short rod about 1.5 μm in length and the other is a longer rod about 4.0 μm in length. Overlaid images of DAPI stained cells showed a compact dense chromosome area (DNA, stained by DAPI) in the cell, while the ribosome FISH signal extended throughout the cytoplasm. In the short rod cells, the chromosome was always located in the middle of the cell, whereas it was closer to one end in longer cells. Our first instinct was that the differences in morphology were related to the presence of two closely related strains of ORM2 (ORM2a and ORM2b) previously identified in the culture from metagenomics (1). To test this hypothesis, specific 16S rRNA qPCR primer sets distinguishing ORM2a and ORM2b (Table S3) were used to quantify each strain in the OR-b1A culture as well as in two additional subcultures (DGG0 and DGG4) that were also were visualized using FISH (Figure 3). The relative abundance of ORM2a via qPCR in these three cultures was 36%, 48%, and 18%, respectively, whereas relative abundance of ORM2b was below 1% in all cases (Figure S1). Since two morphologies were readily observed in samples from cultures DGG0 and OR-b1A despite low relative abundance of strain ORM2b indicated that the two different morphologies were not related to this previously reported strain variation. The different morphologies rather may relate to growth stage or other culture conditions leading to cell elongation.

**Figure 2.**
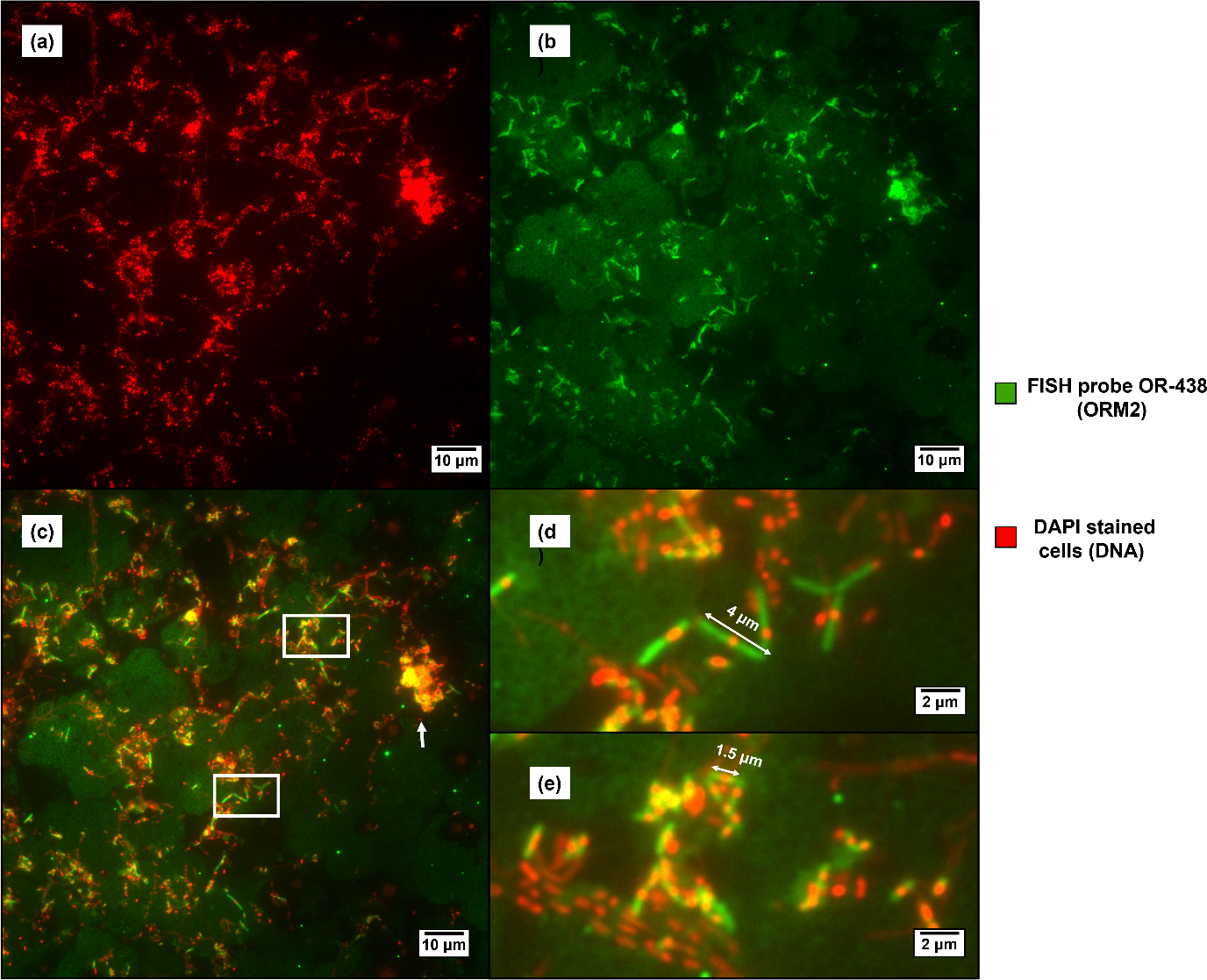
Microscopic images of OR benzene-degrading consortium subculture OR-b1A labelled with FISH probe OR-438 and stained with DAPI. Epifluorescence microscopy images showing only DAPI stained cells (a), cells labelled with probe OR-438 (b), and an overlay image (c) where Desulfobacterota ORM2 cells are in green and all DAPI-stained cells in red. Images are false colored and overlayed using ImageJ. Yellow occurs where a strong OR-438 signal (green) is overlayed with DAPI (red), indicating multiple ORM2 cells. An ORM2 cell aggregate is indicated by a red arrow. Panels (d) and (e) show zoomed in views of the two white rectangles from panel (c).

**Figure 3.**
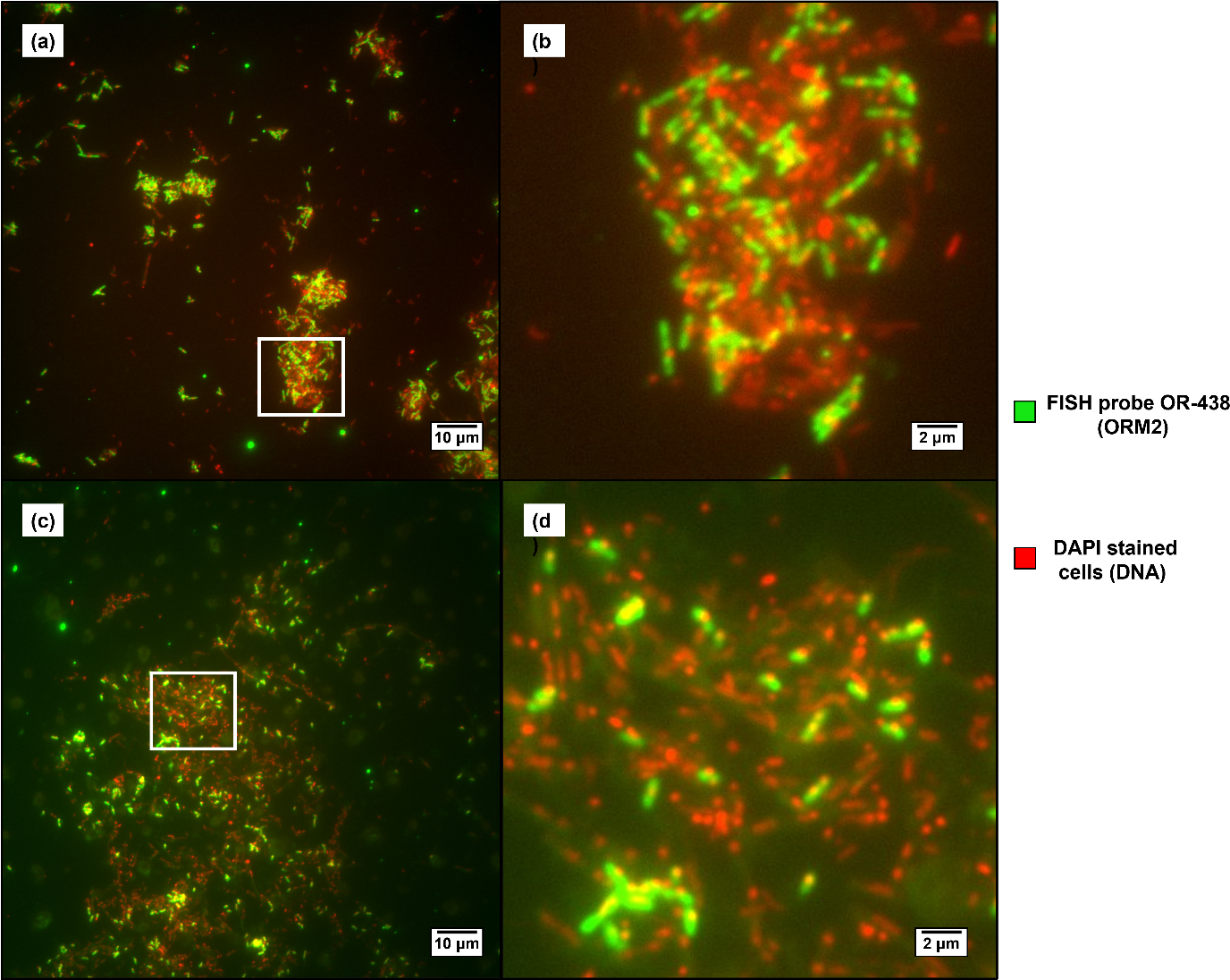
Microscopic images of OR consortium subcultures OR-DGG0 and OR-DGG4 labelled with FISH probe OR-438 labeled and stained with DAPI. Images are false colored and overlayed using ImageJ. Top row (a and b) showing Desulfobacterota ORM2 (green) morphologies in subculture DGG0 and Bottom row (c and d) showing ORM2 morphologies in subculture DGG4. The images to the right (c and d) are zoomed in views of the areas in white rectangles.

### ORM2 co-enriched with *Ca.* Nealsonbacteria, *Ca.* Omnitrophica, *Methanothrix* and *Methanoregula*

To learn more about the morphology and spatial distribution of ORM2 cells within the OR consortium, we selected one subculture (DGG100 original) for a time course growth experiment. Two bottles of sterile, anaerobic minimal medium were inoculated with culture (20% v/v) and maintained for 371 days, reamending benzene whenever concentrations dropped below ∼5 mg/L (∼10 µmol/bottle). The heatmap in Figure 4 provides a comprehensive view of the changes in the absolute abundance of most abundant ASVs over this time. Two ASVs closely matching to ORM2 sequences (annotated as candidate clade Sva0845 Deltaproteobacteria by version 128 of the Silva SSU database, see Table S10) increased in absolute abundance of over time. These two ASVs differ by only one base pair out of 471 nt and have been observed in recent studies (2) but these variants are different from previously identified strains ORM2a and ORM2b (1) from several years ago. The ASVs whose absolute abundances coincided with both ORM2 ASVs were affiliated with two methanogens (*Methanothrix* [formerly *Methanosaeta*] and *Methanoregula*) and two candidate phyla (*Candidatus* Omnitrophica [formerly OP3] and *Candidatus* Nealsonbacteria [formerly OD1]). The co-enrichment of acetoclastic methanogens (*Methanothrix*) and hydrogenotrophic methanogens (*Methanoregula*) is consistent with the production of acetate and hydrogen from benzene fermentation. Moreover, we recently determined that *Ca.* Nealsonbacteria is an ultra-small (∼0.2 µm) epibiont associated with *Methanothrix* and is predicted to be involved in dead biomass recycling (17). As expected, the corresponding ASV clustered with its host *Methanothrix*. *Ca.* Nealsonbacteria’s genomic analysis suggested its potential ability to degrade polysaccharides, which are the main component of extracellular polymeric substances (EPS) (17). *Ca.* Omnitrophica (OP3) belong to the superphylum *Planctomycetes/Verrucomicrobia/Chlamydiae* (PVC) and have been found in many anoxic environments including freshwater sediments (24), wetlands (25), flooded paddy soil (26), wastewater treatment plants and methanogenic bioreactors (27–29). We have not yet analyzed the genome of *Ca.* Omnitrophica, but other genomic studies have proposed that members of this candidate phylum are capable of sulfur cycling and anaerobic respiration (26, 30). In 2022, Kizina et al. (28) successfully visualized a member of *Ca.* Omnitrophica (OP3 Lim) in a methanogenic limonene-degrading cultures using gradient centrifugation and FISH. They found that OP3 Lim were round-shaped ultramicrobacteria 0.2 to 0.3 µm in diameter. Their FISH images revealed OP3 Lim was an epibiont of both archaea and bacteria, though most were found attached to the acetolactic *Methanothrix.* A subsequent proteomics study of the limonene-degrading culture suggested that OP3 Lim was degrading external polysaccharides and/or lipopolysaccharides (28). Although high abundances of *Ca.* Omnitrophica have only been identified in a few of our benzene-degrading subcultures (1), it shares a surprisingly similar lifestyle with what we found for *Ca.* Nealsonbacteria (17). Perhaps they are direct competitors for dead biomass and/or for EPS in our culture. In the experiments conducted in this study, the inoculum culture bottle “DGG100 Original” contained ORM2 and *Ca.* Omnitrophica in similar relative abundances, but fewer *Ca.* Nealsonbacteria compared to other lineages, which may indicate a competitive victory for *Ca.* Omnitrophica in this specific bottle.

**Figure 4.**
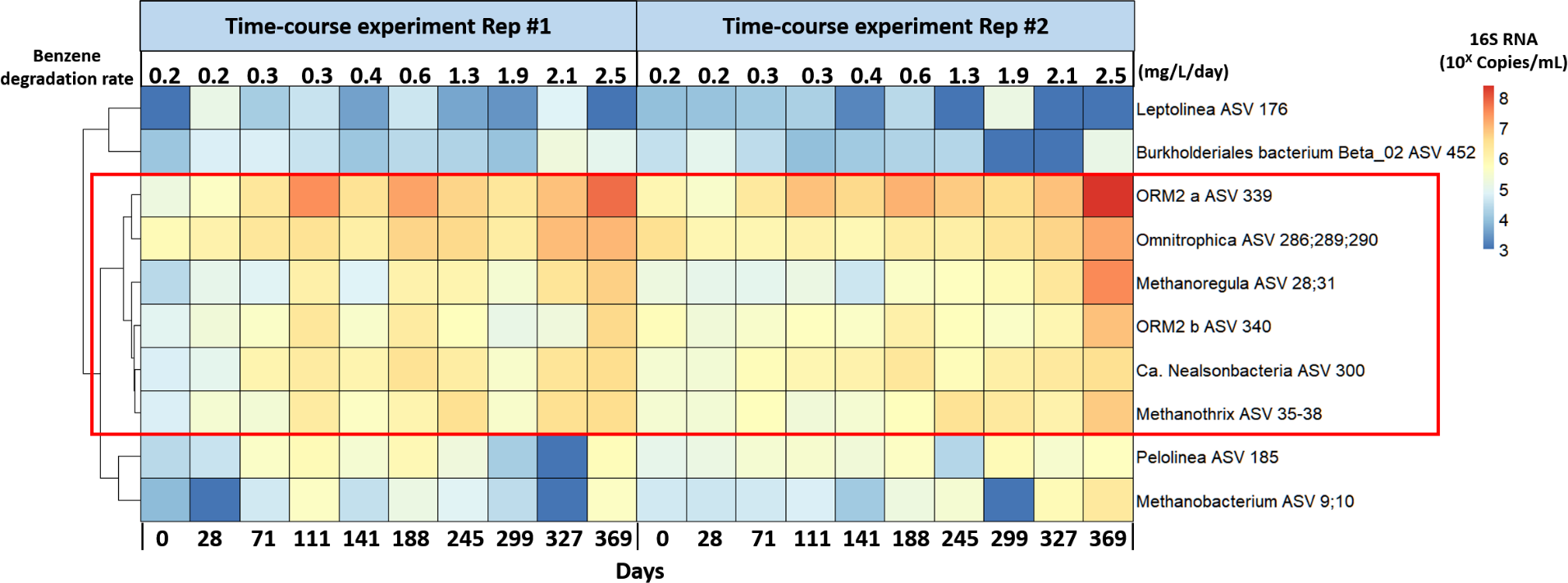
Heatmap of absolute abundance of top ASVs in the time course growth assay. ASVs with relative abundance over 3% were included. Absolute abundance for each ASV was calculated using relative abundance multiplied by absolute total bacteria or archaea determined by qPCR. Numbers at the top of each column represent the benzene degradation rate in mg/l/day. Highly similar ASVs were grouped in one row and their assigned numbers as per Table S10 are indicated.

### The growth of ORM2 seems to occur in phases

Benzene degradation and methane production began without a measurable stall, exhibiting a consistent behavior between duplicate bottles for the full duration of the time course experiment (Figure 5, Top panel). Starting ORM2 concentrations (day 0) in replicates #1 and #2 were 1.7×10^6^ copies/mL and 4.3×10^6^ copies/mL, respectively (Table S6), close to the estimated threshold concentration of 4.3×10^6^ copies/mL required for measurable benzene degradation (1, 2). The initial degradation rate of ∼0.2 mg/L/day in both replicates corresponded to about 20% of the degradation rate of 1 mg/L/day observed in the parent culture (DGG100 original). As time progressed, the absolute abundance of ORM2 measured by qPCR revealed four distinct phases of decay and growth (Table 1 and Figure 5), defined by whether absolute abundances of ORM2 were increasing or decreasing. The first phase, referred to herein as the “decay phase”, lasted between days 0-36 of the growth experiment. In this phase, the culture community composition remained relatively stable (as determined by 16S rRNA gene amplicon sequencing, see bottom panels of Figure 5), yet ORM2 concentrations decreased by an order of magnitude. Using calculations shown in Table S5, the first order decay rate of ORM2 in the “decay phase” was estimated to be −4% per day. It was surprising to see such a substantial drop in ORM2 abundance after inoculating cells into fresh growth medium – the same medium our lab has always used to grow this culture. This decrease is far more than can be explained by the initial 20% dilution of the parent culture (Table S6). The reason behind this ORM2 cell loss is quite puzzling. Since ORM2 cells are the only organisms capable of oxidizing benzene in this culture, and under the assumption that all other microbes in this community rely on the products of ORM2 benzene fermentation for their own survival, one might have expected that concentrations of all other OR microorganisms would also decrease accordingly. Possible explanations for the observed cell decay following culture dilution include physical damage, disruption of cell-cell interactions, shock during transfer, or the dilution of essential enzymes and co-factors (which require extra time to synthesize). Perhaps a large portion of the energy generated from benzene metabolism was partitioned towards non-growth processes, such as producing extracellular enzymes or synthesizing extracellular polymeric substances (EPS) or signaling compounds (31). One might postulate that the decay was simply a result of qPCR error, where DNA from live, dead, and dying cells was amplified. However, a closer inspection of previously published OR cultivation studies reject this hypothesis. Luo et al. (1) conducted a similar cultivation study of the OR consortium but involving much deeper cultures dilutions (1% and 0.1%), and saw near-identical decreases in ORM2 abundances (maximum rate of −3.2% per day) in the first 56 days (Table S7). Guo et al. (6) reported ORM2 decay rates of up to −7% per day in bottles exposed to oxygen (Table 1). These decay rates are remarkably similar.

**Figure 5.**
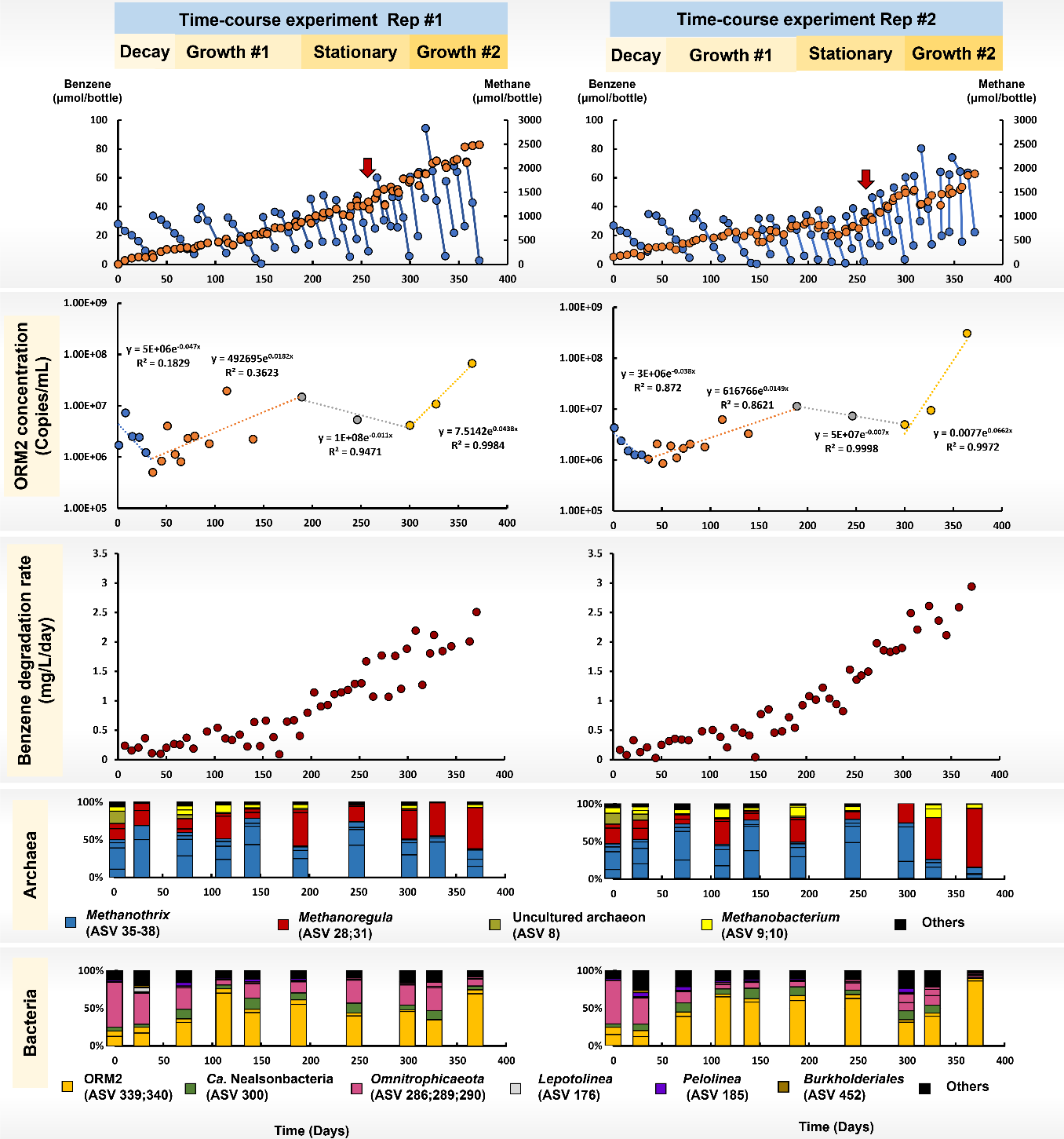
Benzene degradation profile (top panels), ORM2 concentration (second panel), benzene degradation rate (third panel) and relative abundance of top ASVs (bottom two panels) for the time-course experiment. Only ASVs with relative abundance over 3% at least in one of the samples are shown while remaining ASVs are combined as others. The associated 16S amplicon sequencing data are reported in Table S10. Red arrows indicate the time when we began increasing the targe benzene concentration.

**Table 1.** Estimated first-order growth and decay rates and yield for four distinct phases in a time-course growth assay.

| Phases |  | ORM2 | *Ca.*<br>Nealsonbacteria | Bacteria | Archaea |
| --- | --- | --- | --- | --- | --- |
| Decay phase | First order decay rate (day<sup>-1</sup>) | -3.8% to -4.7% | -1.0% to -1.8% | -2% to -2.7% | -0.5% to -1.6% |
| Growth #1 | First order growth rate (day<sup>-1</sup>) | 1.5% to 1.8% | 0.8% to 1.2% | 1.5% to 1.9% | 0.6% to 1.4% |
|  | Measured net yield (g cell/g benzene) | 0.006 to 0.008 | 0.001 | 0.022 to 0.046 | 0.002 to 0.007 |
| Stationary | First order decay rate (day<sup>-1</sup>) | -0.7% to -1.1% | -1.0% to 0.4% | -0.5% to -1.6% | -1% to 0.8% |
| Growth #2 | First order growth rate (day<sup>-1</sup>) | 4.4% to 6.6% | 2.3% to 3.4% | 4.3% to 5.0% | 4.3% to 4.4% |
|  | Measured net yield (g cell/g benzene) | 0.016 to 0.060 | 0.002 | 0.049 to 0.121 | 0.014 to 0.043 |
| Previously Reported Values |  |  |  |  |  |
| Luo et al. 2016 | Growth rate | 2% | NA | NA | NA |
| Luo et al. 2016 | Net yield (g cell/g benzene) | 0.006 to 0.012 | 0.001 to 0.002 | 0.015 to 0.058 | 0.011 to 0.042 |
| Guo et al. 2022 | Growth rate (day<sup>-1</sup>) | 1% to 3% | NA | NA | -0.8% to 2% |
| Guo et al. 2022 | Decay rate (day<sup>-1</sup>) | -2% to -7% | NA | NA | -0.4% to 4% |
| Guo et al. 2022 | Net yield (g cell/g benzene) | 0.02 | NA | NA | 0.019 |

\*Detailed calculations are shown in Table S5. Data shown are ranges from experimental duplicate cultures.

The second phase, “growth phase #1”, commenced on day 36 when ORM2 concentrations began to increase from 5.05×10^5^ copies/mL up to a maximum of 1.96×10^7^ copies/mL by day 189. Rates of anaerobic benzene degradation also increased with ORM2 growth, from 0.1 mg/L/day up to 0.9 mg/L/day. The relative abundance of ORM2 also increased, comprising more than half of all bacteria in the experimental duplicates (Figure 5, bottom panel). As seen in Table 1, the net yield of ORM2 in “growth phase #1” ranged from 0.006 to 0.008 g cell dry mass/g benzene with an average growth rate of approximately 2% per day; these estimates are nearly identical to values previously reported by Luo et al. (1).

Beginning on day 189, the cultures entered a 110-day “stationary phase”, during which ORM2 concentrations plateaued and may have decreased slightly (see row 2 panels in Figure 5). Curiously, rates of anaerobic benzene degradation continued to increase, up to 1.6 mg/L/day (see row 3 panel in Figure 5). Here, we define “stationary phase” as when the rate of cell growth is equal to the rate of cell death/decay. While it has been previously established that ORM2 concentrations correlate to rates of benzene degradation activity in the OR/DGG-B consortium (1, 2), results from this study indicate that the relationship between ORM2 abundances and benzene degradation activity is not perfectly linear and likely one or more auxiliary microbial group in these cultures must be supporting ORM2 activity (20, 32). Methanogens (*Methanothrix* and *Methanoregula*) are clearly vital for degrading benzene fermentation products, but perhaps species associated with dead biomass recycling also help sustain/improve benzene degradation activity. During the “stationary phase”, three ASVs affiliated with *Ca.* Omnitrophica were enriched over time, supporting their predicted role in dead biomass and/or EPS metabolism (Figure 5, bottom panels).

Interestingly, the absolute abundance of ORM2 cells in the “stationary phase” were approximately equal to that of the parent culture (4.85×10^7^ copies/mL). Given that the parent culture and transferred experimental duplicates were amended with similar concentration of benzene (∼20 mg/L), these ORM2 concentrations likely represent the carrying capacity or maximum number of ORM2 cells that can be supported by this substrate concentration. To verify this and to evaluate whether higher concentrations of benzene could support more ORM2 growth and faster benzene degradation activity, we gradually increased the amount of benzene fed to the duplicate cultures from ∼40 µmol/bottle (15 mg/L) up to 80 µmol/bottle (40 mg/L), from day 257 (indicated with red arrows in top panel of Figure 5) through to the end of the experiment. This initiated a second ORM2 growth phase, “growth phase #2”, shortly after Day 300, with faster ORM2 growth rates (6% per day) than seen in “growth phase #1” (2% per day). These data suggest that the growth of ORM2 was constrained by the limited availability of benzene during the “stationary phase”. Further, we recovered the highest relative abundance ORM2 ever recorded in the OR consortium’s history – 86% of total bacterial amplicon sequencing reads (day 369, replicate #2).

### Microscopic images reveal cell aggregation

To study the spatial organization of bacterial and archaeal cells during these different growth phases, samples from the two transfer cultures were analyzed by microscopy, stained using DAPI and FISH probes. First, we compared the parent culture and a sample of the transfer culture during the “decay phase” (Figure 6, first two left images). The parent culture had many large aggregates of cells > 40 µm in diameter, whereas only small aggregates (< 10 µm) could be seen in the diluted culture on day 14 “decay phase”. Note that the bright yellow areas in the images occur where there is significant overlay of the ORM2 probe signal (green colour) on top of the DAPI signal (red colour), corresponding to a high density of ORM2 and indicative of ORM2 cell aggregates. A series of images showing separate channels for the ORM2 probe OR-438, the Archaea probe Arch-915, DAPI, and an overlay of all three channels is provided in Figure S3. These data strongly indicate that physical disruption of larger aggregates occurred during the sampling, transferring, and/or dilution of the parent culture into fresh medium. The cell aggregates, which contain EPS, serve as potential storage for secondary metabolites, quorum-sensing signaling molecules, cell debris, and perhaps essential cofactors (33). When performing culture dilutions, the act of sampling and mixing may disrupt the structure of these cell aggregates, requiring a longer recovery time. Moreover, the introduction of fresh growth medium may trigger the diffusion of stored nutrients from the cell aggregates, potentially slowing down the growth of ORM2 (34). The re-establishment of a stable microenvironment for cell aggregates may therefore be a crucial factor contributing to the observed “decay phase” following inoculation.

**Figure 6.**
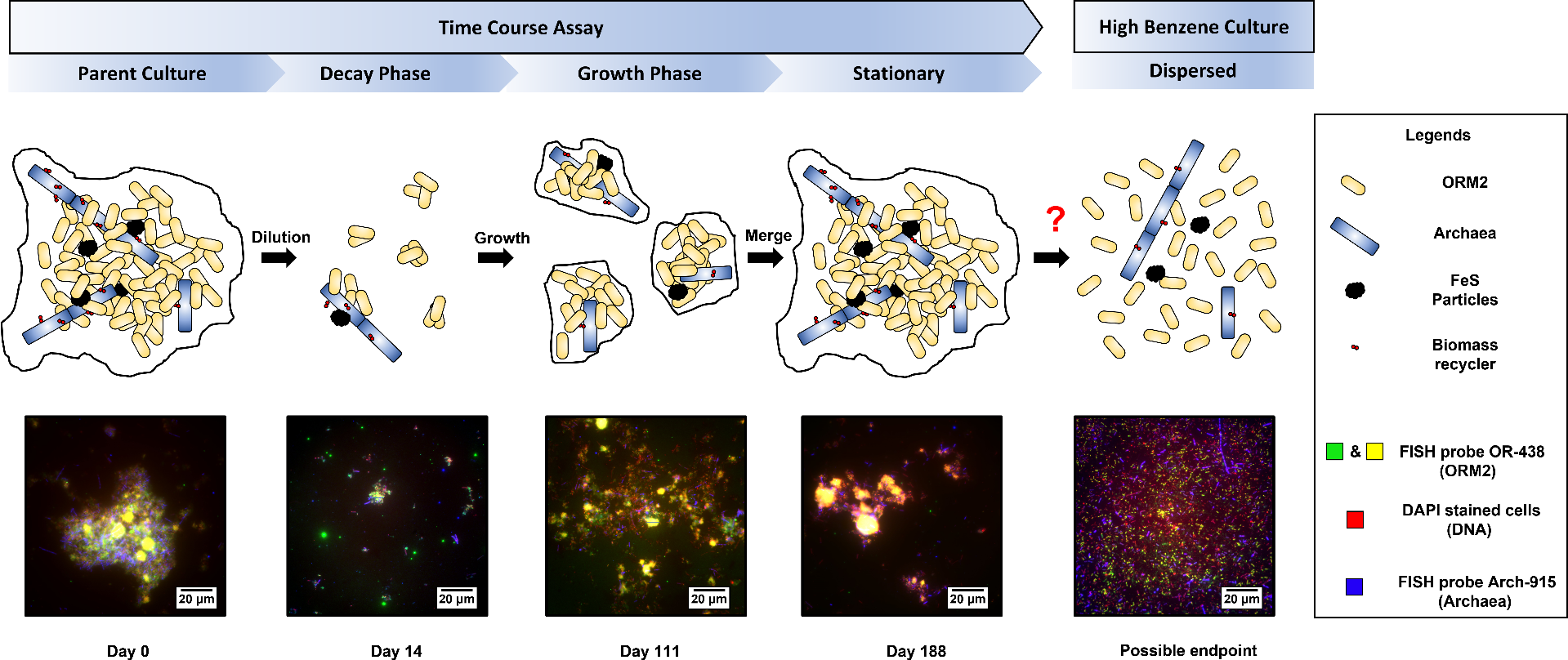
A sketch (top row) of the growth of ORM2 cell aggregates and the related FISH images (bottom row) The date shown at the bottom of the FISH images indicate the sampling time for each image. “Possible endpoint” shows a subsequent sub-transfer culture maintained at high benzene concentration (120 mg/l) which is used a proxy for the endpoint of the time-course growth assay after gradually increasing the benzene feeding amount. Epifluorescence microscopy showing archaea (blue), DAPI (red) and Desulfobacterota ORM2 (Green and yellow). Yellow areas result from overlay of ORM2 probe signal and DAPI signal, indicative of ORM2 cell aggregates.

In “growth phase #1”, both the number and the size of cell aggregates increased (Figure 6, day 111). We identified ORM2 in these cell aggregates as well as methanogenic archaea, iron sulfide particles (a reducing agent in the culture growth medium), and other microbial partners. The cell aggregates provide direct contact between ORM2 and methanogenic archaea which may facilitate exchange of benzene fermentation products (H_2_/CO_2_ and acetate), secondary metabolites or possibly electrons (35) (direct interspecies electron transfer, DIET). In the “stationary phase”, the spatial organization of the duplicate transfers resembled that of the parent culture, comprised primarily of large clusters of ORM2, methanogens, and other microbes (Figure 6 day 188). Typically, cell-aggregated structures are held together with EPS. Producing EPS requires a significant amount of energy consumption and forming cell aggregates and usually results in a motility loss which prevents microbes from moving towards nutrient-rich areas (36, 37). Although forming cell aggregates is energetically costly, it creates a stable environment for microbes to grow. These microscopic images clearly show that the cell aggregates are primarily composed of ORM2 cells, particularly when the cell concentration is very low. Other researchers have found that this kind of cell aggregation usually occurs under substrate-transfer-limited conditions, where the local consumption of substrate creates substrate gradients that help the cells in aggregates obtain more substrate (38–40).

Regrettably, as a result of the COVID-19 pandemic lockdown, we were unable to collect microscopic samples for “growth phase #2”. Only DNA samples were obtained and analyzed during this period. In order to gain insight into how the culture would behave if we continued to increase the liquid benzene concentration as we did in “growth phase #2”, we imaged another culture, “DGG100 PosB”, which is a 100% transfer of the time-course assay parent culture “DGG-100 original” (Figure S1). The establishment of “DGG100 PosB” was part of an oxygen tolerance experiment conducted by Guo et al. (6), where it served as a strictly anaerobic positive control designated as Bottle 5. In the study by Guo et al., the “DGG100 PosB” culture was incubated for 422 days, during which the concentrations of benzene amended at each feeding gradually increased from ∼10 mg/L to 159 mg/L (Table S8 and Figure S4). The “DGG100 PosB” shared the same parent culture as our time-course assay and exhibits similar behavior as benzene feeding concentration increased over time. Therefore, we selected the “DGG100 PosB” as a representative example for the endpoint for the time-course growth experiment, as we continued to increase benzene concentration. The FISH images for the “DGG100 PosB” culture were obtained on the day 308 of the experiment conducted by Guo et al. (6) with an approximate benzene feeding concentration of 102 mg/L. These images illustrate the spatial distribution of the methanogenic benzene-degrading culture under high benzene liquid concentration conditions.

### ORM2 cell aggregates are likely responding to changes in benzene concentration

At the time of sampling, the “DGG100 PosB” culture was actively degrading benzene at a rate of 5 mg/L/day, or roughly 4 times faster than the “DGG100 original” culture (1.3 mg/L/day). When ORM2 concentrations were quantified in each bottle, we discovered that their abundances were near-identical (2.25×10^7^ cells/mL for DGG100 PosB and 4.85×10^7^ cells/mL for DGG100 original), as was also seen in the experimental bottles in the “stationary phase”. However, when we looked at the cells under the microscope, we observed that the spatial distribution was completely different. We found that *Desulfobacterota* ORM2 and methanogenic archaea were evenly distributed and dispersed throughout the culture, as shown in Figure 6 (labelled “possible endpoint”), in stark contrast to large cell aggregates previously seen. Each of these cultures was maintained in a near-identical manner – the only difference was that DGG100 PosB was fed 3 – 10 times more benzene than the other bottles. The absence of cell clusters in DGG100 PosB indicates that aggregation may be a response to low benzene concentrations (< 40 mg/L). EPS consists mainly of polysaccharides and proteins which cover and bind cells as sheaths or in slime (33, 37, 41). EPS can modulate cell surface charge and increase hydrophobicity (42) by virtue of the composition and abundance of proteins and lipids, and thus may increase the local benzene concentration in the surrounding environment of cell aggregates via diffusion or absorption (33, 37). Yang et al. (43) measured higher benzene concentrations in the EPS of biosludge which may be similar to our ORM2 cell aggregates. When bulk benzene aqueous concentrations are low, the higher local concentrations afforded by cell aggregation and EPS production could benefit the growth of ORM2. However, when the bulk benzene aqueous concentration is much higher, the EPS could have the reverse effect owing to unfavorable osmotic pressure (44) and heightened toxicity of benzene, both of which can inhibit the production of EPS and the growth of microbes. Perhaps higher substrate (i.e. benzene) concentrations cause down-regulation of EPS-producing genes as a defense mechanism against the toxicity of high benzene concentration, as reported elsewhere (42), partitioning more energy for ORM2 cell growth (37). Moreover, the EPS released from cell aggregates could be utilized by other EPS degraders in the culture as their carbon and energy source. The consumption of EPS would expectedly break the ORM2 cell aggregates and result in more free-living ORM2 cells. Compared to the cell aggregates, free-living ORM2 cells have a higher surface-to-volume ratio which may also promote benzene degradation and result in the higher benzene degradation rates seen in the DGG PosB culture. This hypothesis requires further testing with a more detailed transcriptome/proteome study and metabolites analysis of the cell aggregates.

### ORM2 genome and proteome analysis reveals processes related to EPS assembly and export

The complete genome of *Desulfobacterota* ORM2 was examined for genes related to EPS assembly and export. Since the genes for monosaccharide production are not specific for EPS, we searched for more conserved genes for the subsequent steps of polymerization, assembly, and export (45, 46). Although the understanding of EPS production is limited, there are three primary mechanisms that have been described in literature, including Wzy-(O-antigen polymerase-), ABC transporter- and synthase-dependent pathways (45, 47). The ORM2 closed genome was examined, and 26 genes were found related to polysaccharide/lipopolysaccharide/ exopolysaccharide synthesis, assembly, and export, of which 6 were expressed in a parallel proteomics study (Table S9; (23)). Moreover, 34 genes were found related to ABC transporters and 16 genes were annotated as tetratricopeptide repeat protein (TPR) which are involved in synthase-dependent pathways (48). Three genes affiliated to ABC transporter pathway and two TPR genes were found expressed in previous proteomics studies (Table S9) (23). The existence and expression of EPS-related genes suggest that ORM2 is capable of producing EPS, consistent with our microscopic observations.

Genes related to EPS production were also examined in the entire (OR consortium) metagenome. Genomic bins belong to *Ca.* Nealsonbacteria, *Spirochaetota*, *Desulfobacterota*, *Bacteroidales*, *Syntrophales*, *Ca.* Omnitrophica, *Methanoregulaceae*, and *Methanothrix* were investigated (Table S9). Bins from methanogens contained very few genes related to EPS production, suggesting that they do not participate in EPS production. Conversely, most of bacterial bins examined contained numerous genes involved in lipopolysaccharide/polysaccharide synthesis and export – but were not detected by proteomics (23). It is conceivable that these bacteria do possess the ability to produce EPS, but due to the low abundance of these microbes in the OR/DGG-B consortium, the corresponding proteins are undetected and their contribution to EPS and cell aggregation likely minimal.

### Implications

In this study, a FISH probe specific to a tight group of benzene-fermenting organisms represented by Desulfobacterota ORM2 was designed, validated, and used to successfully visualize the morphology of these cells. This new probe provides a means of monitoring the appearance of the key benzene-degrader ORM2 and other major phylotypes in this methanogenic benzene degrading culture. Microscopy combined with a long term growth experiment revealed that the loss of ORM2 cells after inoculation may be due to disruption and reassembly of cell aggregates requiring energy for the synthesis of EPS. The EPS maintains the structure of cell aggregates, and its hydrophobicity may increase the local concentration of benzene, thereby promoting growth at low benzene concentrations. Such aggregates may facilitate direct contact between the benzene fermenter ORM2, acetoclastic and hydrogenotrophic methanogens, biomass and EPS recyclers, and FeS particles used to reduce the medium. This close association enhances the nutrient exchange and the transfer of secondary metabolites, hydrogen, acetate, co-factors and may promote potential direct interspecies electron transfer (DIET). Liu et al. reported that the DIET could potentially take place via EPS (35). Furthermore, the trapped FeS particles may also be used as a conductive matter to promote DIET (49). The cell aggregates also retain extracellular enzymes and other exudates, vesicles, nutrients, vitamins, and quorum-sensing molecules, promoting a stable and sustainable environment for the entire community. The ORM2 genome also encoded many genes related to EPS production and export. Finally, cell aggregation may be related to the prevailing aqueous benzene concentration, although confirmatory studies are required. A high benzene liquid concentration may sustain higher rates of growth while the nutrient concentrations precluding the need for cell aggregation. The ability to visualize benzene-fermenting *Desulfobacterota* ORM2 cells within the associated methanogenic community now provides a new perspective for understanding interspecies interactions and could potentially help improve growth rates and cultivation challenges for this and similar cultures.

### Supporting information

The PDF of a word file contains: 1) supplementary Figures S1 to S3, including a subculturing history of “OR consortium”, FISH images with probe DELTA-495a and a demonstration of different signal channels for FISH images; 2) supporting texts (S1) providing additional description of maximum likelihood phylogenetic tree construction; 3) supplementary tables S3, S6 and S7 listing primers, qPCR results for OR-b1A and calculated half-life and decay rates. The supporting excel file contains larger data Tables S1-S2, S4-S5 and S8-S9.

## Supporting information

Supplemental large tables

## Acknowledgements

We gratefully acknowledge the funding provided by the Genomic Application Partnership Program (GAPP) through grants awarded to E.A.E. (Project IDs OGI-102 and OGI-173). This research was made possible by the generous support of Genome Canada, Ontario Genomics, the Government of Ontario, Mitacs Canada, SiREM, Alberta Innovates, Federated Co-operatives Limited, and Imperial Oil. Special thanks are extended to SiREM (Guelph, Ontario N1G 3Z2, Canada) for providing the culture used in our experiments. We would also like to express our appreciation to Dr. Neil Thomson (University of Waterloo), Dr. Ania Ulrich (University of Alberta), SiREM, and Innotech Alberta (Edmonton, Alberta T6N 1E4, Canada) for their valuable feedback and intellectual contributions during weekly GAPP consortium meetings, which greatly enriched this study.

## Notes

### Competing Interest Statement

The authors have declared no competing interest.

