## Supplemental information for "Visualization of syntrophic benzene-fermenting *Desulfobacterota* ORM2 in a methanogenic enrichment culture using fluorescence *in situ* hybridization"

Number of Pages: 9

Number of SI Figures: 4
Number of Supporting Texts: 1
Number of SI Tables: 10

**Table of Contents**

**Text S1: Description of maximum likelihood phylogenetic tree (Figure 1) construction.**

**Figure S1. Subculturing history of "OR consortium".**
**Figure S2. Microscopic images of FISH probe DELTA-495a labeled OR consortium subcultures – OR-b1A**.
**Figure S3. Demonstration of different signal channels and an overlayed image**.
**Figure S4. Benzene degradation profile of culture DGG100 PosB**.

**Table S1: Summary of benzene and methane concentration (mg/L) and their calculated amounts (µmol/bottle) in time course experiment (Excel file)**
**Table S2. Information on FISH probes used in this study (Excel file)**
**Table S3. Specific 16S rRNA primers used for qPCR in this study.**
**Table S4. FISH probe DELTA-495a blast results against DGG0 and DGG1 metagenome (Excel file)**
**Table S5. qPCR for time course FISH assay and calculated growth/decay rates and doubling times/half-lives (Excel file)**
**Table S6. qPCR results for parent culture (OR-b1A) and the comparison of minimum concentration during lag phase.**
**Table S7. Half-life and decay rate calculated based on experimental data published by Luo et al (1).**
**Table S8. Summary of benzene and methane concentration (mg/L) and their calculated amounts (µmol/bottle) in culture "DGG100 PosB"(Excel file)**
**Table S9. List of all ORM2 genes related to EPS production (Excel file)**
**Table S10: List of all gDNA samples and 16S rRNA gene amplicon sequencing results for time-course assay (including taxonomic assignment and sequences) (Excel file)**

**References for SI**

**Text S1: Description of maximum likelihood phylogenetic tree (Figure 1) construction.**
MAFFT were used to align full-length and partial 16S rRNA gene sequences in Geneious version 8.1.9. Sites with over 30% gaps were removed. The maximum likelihood tree was constructed based on the alignment file with RAxML 1.0 in Geneiuous version 8.1.9 with GTRGAMMA model and 100 bootstrap replicates. The tree was then edited with Mega11. Please note that KT322260.1, EF613404.1, AB671643.1, JQ367316.2, EF613434.1, KJ635794.1, JF323734.1 and Syntrophorhabdaceae UBA5609 were partial 16S rRNA gene sequences less than 1000 base pairs.

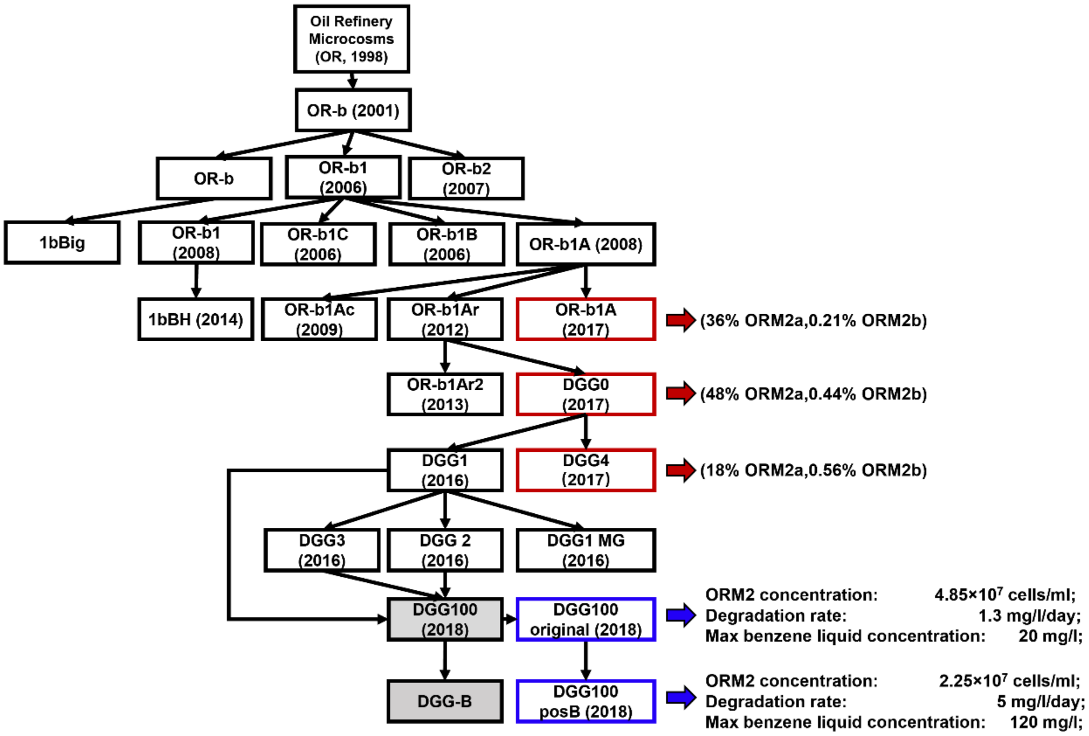


**Figure S1. Subculturing history of "OR consortium".** Each name corresponds to a subculture and the dates indicate the year when the subculture was first established. The red rectangles indicate subcultures used for ORM2 morphology examination and the blue rectangle indicates the subcultures used for time-course FISH experiment. Boxes with grey shade are bigger vessels (>100 L) maintained in SiREM labs.

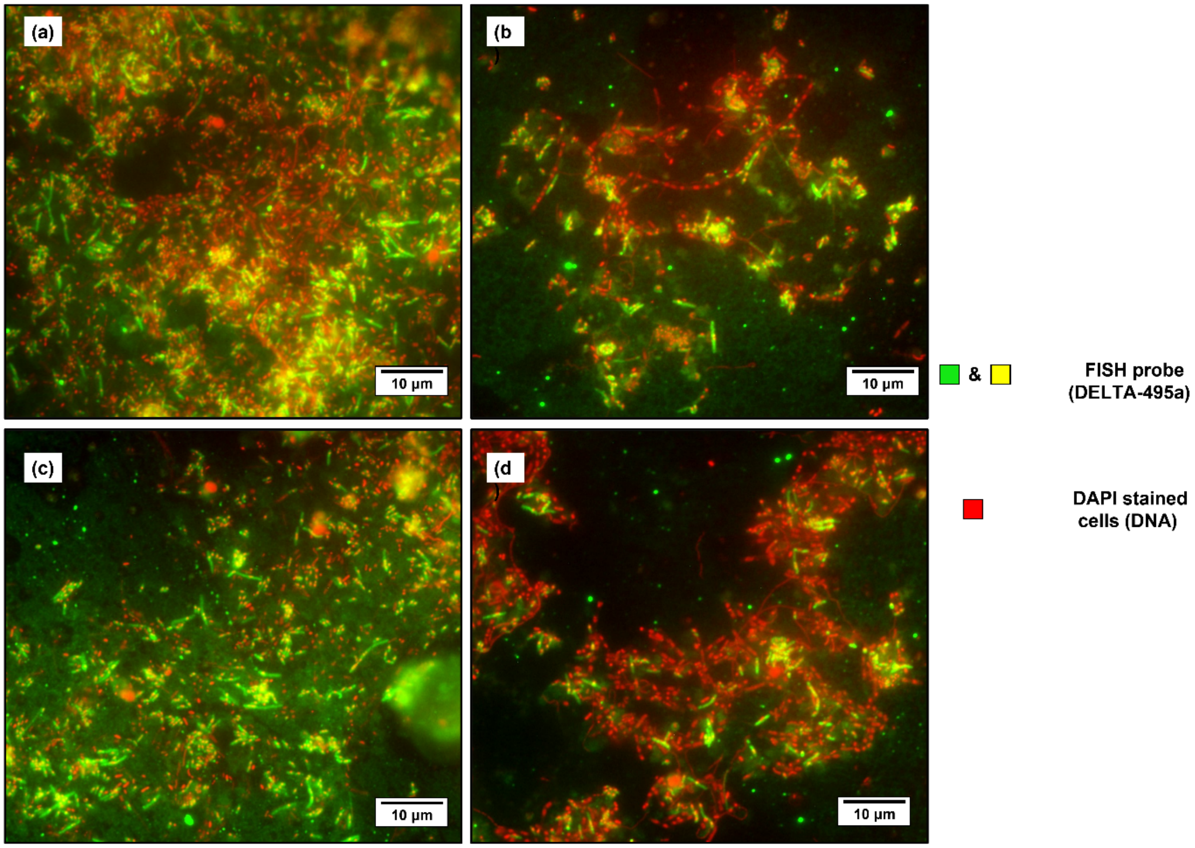


**Figure S2. Microscopic images of OR consortium –OR-b1A – labelled with FISH probe DELTA-495a and stained with DAPI**. Panels (a-d) show representative FISH images with FISH probe DELTA-495a and DAPI. Images are false colored and overlayed using ImageJ. Epifluorescence microscopy DAPI (red) and DELTA-495a (Green and yellow). Yellow areas come from the overlay of DELTA0495a probe signal and DAPI signal (see Figure S3).

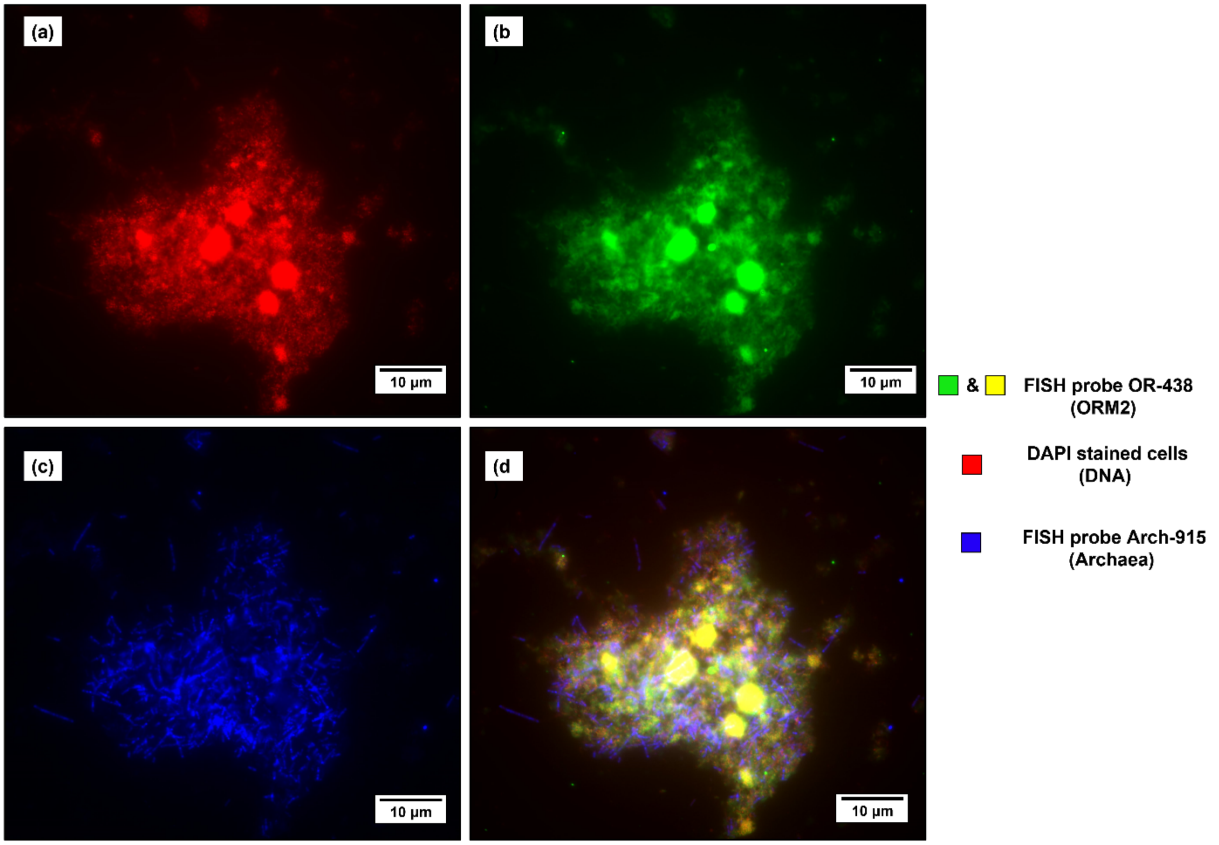


**Figure S3. Demonstration of different signal channels and an overlayed image of Parent culture DGG100 original**. a) DAPI stained signal; b) FISH probe OR-438 signal; c) FISH probe Arch-915 signal and d) overlayed image. Images are false colored and overlayed using ImageJ. Epifluorescence microscopy showing archaea (blue), DAPI (red) and ORM2 (green and yellow). Yellow comes from overlay of green ORM2 FISH probe signal over red DAPI signal, and is indicative of ORM2 cell aggregates.

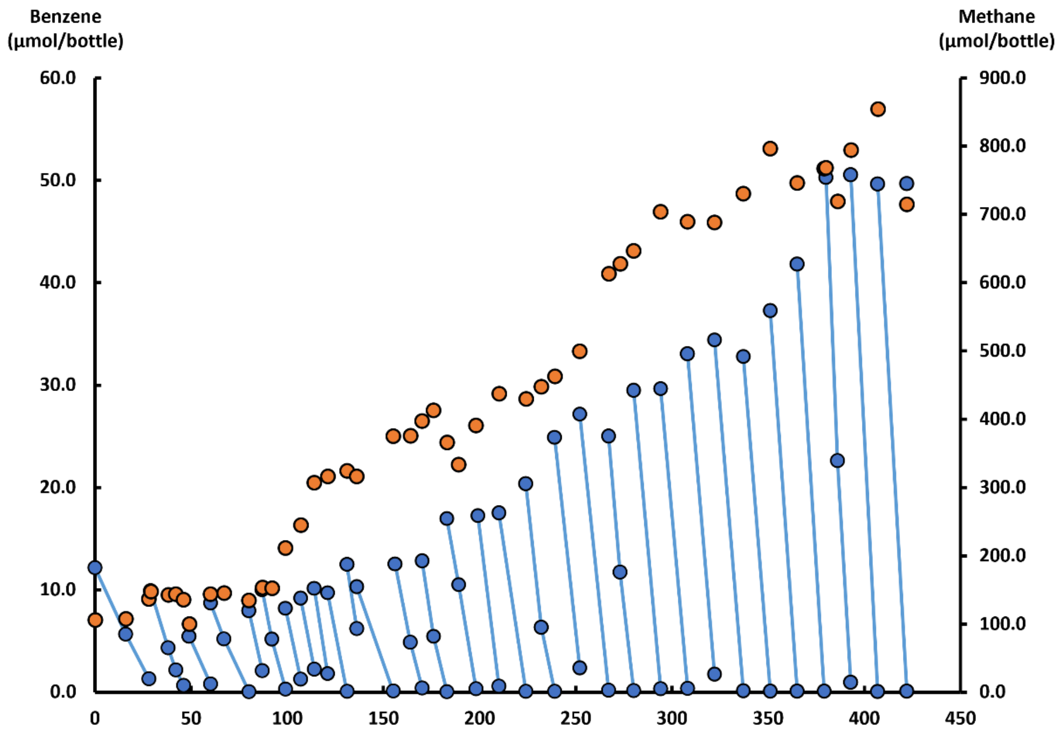


**Figure S4. Benzene degradation profile of culture DGG100 PosB**. Blue dots represent the benzene concentration, while orange dots represent the methane concentration. The red arrow indicates the time of sampling for FISH imaging. This data was reported in Guo et al., 2022.

**Tables S1-S2, S4-S5 and S8-S10** are larger and are provided in the associated Excel file.

**Table S3. Specific 16S rRNA primers used for qPCR in this study.**

| Primer Name | Target Organism | Primer Sequence (5' - 3') | Expected Amplicon Length | Primer Set Specificity* | Annealing Temperature (°C) | References |
|---|---|---|---|---|---|---|
| Bac_1055f | General Bacteria | ATGGCTGTCGTCAGCT | 338 | 1085282/3195888 (kingdom Bacteria) | 55 | (Amann et al., 1995, Ferris et al., 1996) |
| Bac_1392r | | ACGGGCGGTGTGTAC | | | | |
| Arch_787f | General Archaea | ATTAGATACCCGBGTAGTCC | 273 | 56410/160768 (kingdom Archaea) | 59 | (Yu et al., 2005) |
| Arch_1059r | | GCCATGCACCWCCTCT | | | | |
| ORM2_168f | Candidate Sva0485 Deltaproteobacterium, ORM2 | GAGGGAATAGCCAAAGGTGA | 274 | 15/3195888 (kingdom Bacteria) | 59 | (Qiao et al., 2018), (Toth et al.,2021) |
| ORM2_422r | | GAGCTTTACGACCCGAAGAC | | 16/14443 (unclassified Deltaproteobacteria spp.) | | |
| ORM2a_24f | Candidate Sva0485 Deltaproteobacterium ORM2 strain a | CGAGAAAGTTCCGTTCGCGGAATG | 216 | 33/3195888(kingdom Bacteria) | 70 | (Luo et al.,2015) |
| ORM2a_239r | | ACTAGCTAATGGGCCGCGGAC | | | | |
| ORM2b_73f | Candidate Sva0485 Deltaproteobacterium ORM2 strain b | GCTTGCAGGATGAGTAAAGTG | 212 | 183/3195888 (kingdom Bacteria) | 65 | (Luo et al.,2015) |
| ORM2b_284r | | GCCTTGGTAGGCTTTTACCC | | | | |
| OD1_987f | The dominant Ca. Nealsonbacteria in the OR/DGG consortium | GGTGCTGCATGGTTGTCGTC | 200 | 47501/3195888 (kingdom Bacteria) | 65 | (Luo et al.,2015) |
| OD1_1186r | | GCTGCCCTCTGTAACTGCCA | | | | |

**Table S6. qPCR results for parent culture (DGG100 original) and the comparison of minimum concentration during lag phase.**

| | ORM2 (copies/ml) | GenBac (copies/ml) | GenArch (copies/ml) |
|---|---|---|---|
| **Parent Culture** | 4.85E+07 | 1.74E+08 | 1.56E+07 |
| **20% of Parent (Calculated)** | 9.70E+06 | 3.48E+07 | 3.12E+06 |
| **Initial concentration of Rep #1** | 1.71E+06 | 1.17E+06* | 1.17E+05 |
| **Initial concentration of Rep #2** | 4.32E+06 | 6.20E+06 | 5.74E+05 |
| **Minimum in decay phase (Rep #1)** | 5.05E+05 | 1.04E+06 | 1.17E+05 |
| **Minimum in decay phase (Rep #2)** | 1.04E+06 | 1.70E+06 | 2.03E+05 |

***value seems a little low as it is less than the ORM2 copies. See Table S5 for all qPCR data**

**Table S7. Half-life and decay rate calculated based on experimental ORM2 qPCR data published by Luo et al (2016).**

| | 1% inoculum | | | | 0.1% inoculum | | | |
|---|---|---|---|---|---|---|---|---|
| **Time (day)** | **Rep#1** | **Rep#2** | **Rep#3** | **Average** | **Rep#1** | **Rep#2** | **Rep#3** | **Average** |
| **0** | 6.31E+04 | 5.25E+04 | 9.92E+04 | 7.16E+04 | 9.30E+03 | 2.24E+04 | 8.09E+03 | 1.33E+04 |
| **57** | 8.59E+04 | 3.92E+04 | 3.50E+04 | 5.33E+04 | 1.05E+04 | 3.71E+03 | 2.13E+03 | 5.46E+03 |
| **Half life** | N/A | -135 | -38 | -134 | N/A | -22 | -30 | -44 |
| **Decay rate** | N/A | -0.5% | -1.8% | -0.5% | N/A | -3.2% | -2.4% | -1.6% |
